# Phenylacetic acid metabolism in land plants: novel pathways and metabolites

**DOI:** 10.1101/2024.11.06.622264

**Authors:** Pavel Hladík, Federica Brunoni, Asta Žukauskaitė, Marek Zatloukal, Jakub Bělíček, David Kopečný, Pierre Briozzo, Nathan Ferchaud, Ondřej Novák, Aleš Pěnčík

## Abstract

In recent years, substantial progress has been made in exploring auxin conjugation and metabolism, primarily aiming at indole-3-acetic acid (IAA). However, the metabolic regulation of another key auxin, phenylacetic acid (PAA), remains largely uncharacterized. Here, we provide a comprehensive exploration of PAA metabolism in land plants. Through LC-MS screening across multiple plant species and their organs, we identified four previously unreported endogenous PAA metabolites: phenylacetyl-leucine (PAA-Leu), phenylacetyl-phenylalanine (PAA-Phe), phenylacetyl-valine (PAA-Val), and phenylacetyl-glucose (PAA-glc). Enzyme assays, genetic evidence, crystal structures, and docking studies demonstrate that PAA and IAA share core metabolic machinery, revealing a complex regulatory network that maintains auxin homeostasis. Furthermore, our study of PAA conjugation with amino acids and glucose suggests limited compensatory mechanisms within known conjugation pathways, pointing to the existence of alternative metabolic routes in land plants. These insights advance our knowledge of auxin-specific metabolic networks and highlight the unique complexity within plant hormone regulation.

## Introduction

Auxins are a class of phytohormones that are essential for coordinating plant growth and development. Indole-3-acetic acid (IAA) has been extensively investigated due to its diverse physiological impacts (Davies, 2010). Phenylacetic acid (PAA), another constituent of the auxin family, has recently gained attention due to its potential significance in plant physiology and auxin signalling pathways. Although PAA accumulates more than IAA in most plant species, its homeostasis and function are not yet fully understood (Wightman & Lighty, 1982; Sugawara et al., 2015; Cook, 2019). The auxin activity of PAA has been estimated to be less than 10% of that of IAA by three typical auxin tests: the cylinder test, the oat bending test, and the pea test (Haagen-Smit & Went, 1935). However, its main activity appears to be in lateral root induction and root growth promotion (reviewed in Cook, 2019; Perez et al., 2023). Additionally, PAA has antimicrobial properties, exhibiting anti-fungal and anti-bacterial activities (Kawazu et al., 1996; Liu et al., 2014; Zhang et al., 2022). Plants increase PAA production when attacked by herbivores, and exogenous PAA application has been reported to protect plants against fungal pathogens (Perez et al., 2023). However, the exact biological function of PAA in plant defence mechanisms is still unclear and requires further evidence (Kunkel & Harper, 2018).

Biosynthesis of PAA originates from phenylalanine (Phe) and parallels IAA biosynthesis, albeit with different enzymes. Arogenate dehydratase alters PAA levels in Arabidopsis by converting arogenate to Phe, highlighting a Phe-dependent pathway (Aoi et al., 2020a). The primary route involves converting Phe to phenylpyruvate, which then decarboxylates to PAA, similar to the IAA pathway (Cook et al., 2016). However, enzymes responsible for this conversion, like phenylpyruvate aminotransferase from petunia, are still being studied (Cook et al., 2016; Yoo et al., 2013). Secondary pathways include converting Phe to phenylacetaldehyde (PAAld) *via* phenylacetaldehyde synthase (Kaminaga et al., 2006) or to phenylethylamine by aromatic amino acid decarboxylases (Tieman et al., 2006). Aldehyde dehydrogenase family 2 (ALDH2) might be involved in the oxidation of PAAld to PAA as four maize isoforms were shown to be highly active towards many aromatic aldehydes including PAAld (Končitíková et al., 2015). Additionally, a minor, stress-activated pathway converts Phe to phenylacetaldoxime *via* CYP79A2, which is directly converted to PAA in maize (Perez et al., 2021).

The mechanisms for PAA inactivation employ similar pathways involved in IAA inactivation. The majority of IAA in plants exists in its non-active form, which can be categorised into two main groups. The first comprises reversible storage forms, such as esters and amides, or methylated IAA. The second group consists of oxidised metabolites, which undergo irreversible metabolism leading to their degradation (reviewed in Casanova-Sáez et al., 2021; Cohen & Strader, 2024). In PAA metabolism, only the first group has been described, as no oxidative metabolites had been identified. Earlier work in 2005 revealed that GRETCHEN HAGEN 3 (GH3) proteins, known for their role in forming IAA-amides with amino acids (IAA-AAs), also exhibit *in vitro* sensitivity to PAA (Staswick et al., 2005). Subsequently, the first two conjugates, PAA-aspartate (PAA-Asp) and PAA-glutamate (PAA-Glu), were identified in transgenic *Arabidopsis thaliana* plants expressing β-estradiol-inducible YUCCA enzymes, whose induction led to the increase of endogenous levels of PAA-Glu by 14 to 41-fold and PAA-Asp levels by 1.6 to 3.8-fold (Sugawara et al., 2015). The GH3 enzyme involvement in PAA metabolism *in planta* has been confirmed in several studies. According to Sugawara et al., (2015), induction of *GH3*.*9* in β-estradiol-inducible Arabidopsis *GH3*.*9* transgenic plants resulted in an endogenous PAA-Glu level increase by 13-fold. Arabidopsis *GH3*.*5* overexpressing plants accumulated 15 to 70-fold higher PAA-Asp levels than wild-type plants, while PAA levels decreased by up to 5-fold (Westfall et al., 2016). Moreover, the application of PAA or IAA to wild-type plants reciprocally reduced levels of opposite active auxin by increasing the corresponding aspartate metabolites in a GH3-dependent manner (Aoi et al., 2020b). Other IAA metabolites conjugated with amino acids, such as IAA-Ala, -Gly, -Leu, -Phe, -Trp and -Val, have also been detected in plants (Kowalczyk & Sandberg, 2001; Pěnčík et al., 2009; Staswick, 2009). However, their concentrations are typically much lower than those of IAA-Asp and IAA-Glu, likely due to rapid conversion to free IAA mediated by enzymes such as IAA-LEUCINE RESISTANT 1 (ILR1), ILR1-LIKE proteins (ILLs), and IAA-ALANINE RESISTANT 3 protein (IAR3) (Bartel & Fink, 1995; Davies et al., 1999; LeClere et al., 2002), or through their oxidation to oxIAA-amino acids (oxIAA-AAs) (Hladík et al., 2023). Notably, from these low abundant amino acid conjugates, only PAA-Trp was identified in Arabidopsis at concentrations 17-fold higher than its IAA counterpart, suggesting a potential endogenous role for this PAA metabolite (Staswick et al., 2017).

An alternative pathway in auxin metabolism involves the formation of glucosides, such as IAA or oxIAA glucosyl ester (IAA/oxIAA-glc), catalysed by the enzymes UGT74D1 and UGT84B1 (Jackson et al., 2001; Mateo-Bonmatí et al., 2021). In *in vitro* experiments, both IAA and PAA have been shown to serve as substrates for UGT84B1. However, only IAA-glc has been detected *in vivo* (Aoi et al., 2020c; Grubb et al., 2004). IAA methylation, mediated by the IAA CARBOXYMETHYLTRANSFERASE 1 (IAMT1) enzyme, has been demonstrated in plants (Qin et al., 2005). However, overexpression of the IAMT1 in Arabidopsis did not lead to a reduction in PAA levels, suggesting that this enzyme is not responsible for PAA methylation in plants. Nevertheless, PAA methyl ester has been identified in *Escherichia coli* (E. coli) and therefore its presence in plants cannot be ruled out (Takubo et al., 2020). Regardless of whether PAA has been widely detected in the plant kingdom, mechanisms for its inactivation, which may be shared among other species, have only been investigated in Arabidopsis.

Despite recent discoveries in PAA homeostasis, our understanding of how PAA is metabolized in plants remains incomplete. In this study, we identified PAA glucosyl ester (PAA-glc) *in planta* for the first time, as well as three novel endogenous amino acid conjugates, phenylacetyl-leucine (PAA-Leu), phenylacetyl-phenylalanine (PAA-Phe) and phenylacetyl-valine (PAA-Val). A quantitative profiling of a range of PAA metabolites across a spectrum of model plant species, spanning from Bryophyta to Angiosperms, performed by high-performance liquid chromatography-tandem mass spectrometry (HPLC-MS/MS), revealed differences in PAA metabolism as distribution of conjugates differed notably among the studied species as well as their organs. To elucidate PAA metabolic pathways, we further performed bacterial enzyme assays to identify the candidate enzymes for inactivation/activation of PAA and expanded those finding *in planta* by a feeding assay using PAA in different genetic backgrounds or upon chemical knockdown of the IAA-conjugation pathway.

## Results

### PAA-glc is an endogenous PAA metabolite synthesized by UGT84B1 and UGT74D1 glucosyltransferases

To determine PAA conjugates, we adopted and modified a method previously developed and applied for IAA metabolites profiling (Hladík et al., 2023). Having this dependable analytical method, we systematically screened for PAA conjugates across various species of land plants. Remarkably, we uncovered the presence of endogenous PAA-glc, a compound previously undetected in plants, within three species: Arabidopsis, pea and spruce. To ensure the identity of endogenous PAA-glc, we compared its chromatographic retention times from Arabidopsis and spruce extracts with that of synthetic PAA-glc standard (**Fig. S1**). Subsequently, to further confirm the formation of PAA-glc *in planta*, we treated Arabidopsis seedlings with 20 μM PAA, and PAA-glc levels were subsequently determined after 30, 60, and 180 min intervals (**Fig. 1A**). Notably, the concentration of PAA-glc progressively increased from ≈180 pmol g^-1^ to 40 nmol g^-1^ FW after 180 min of treatment, demonstrating the *de novo* synthesis of PAA-glc in response to exogenous application of PAA.

**Figure 1:**
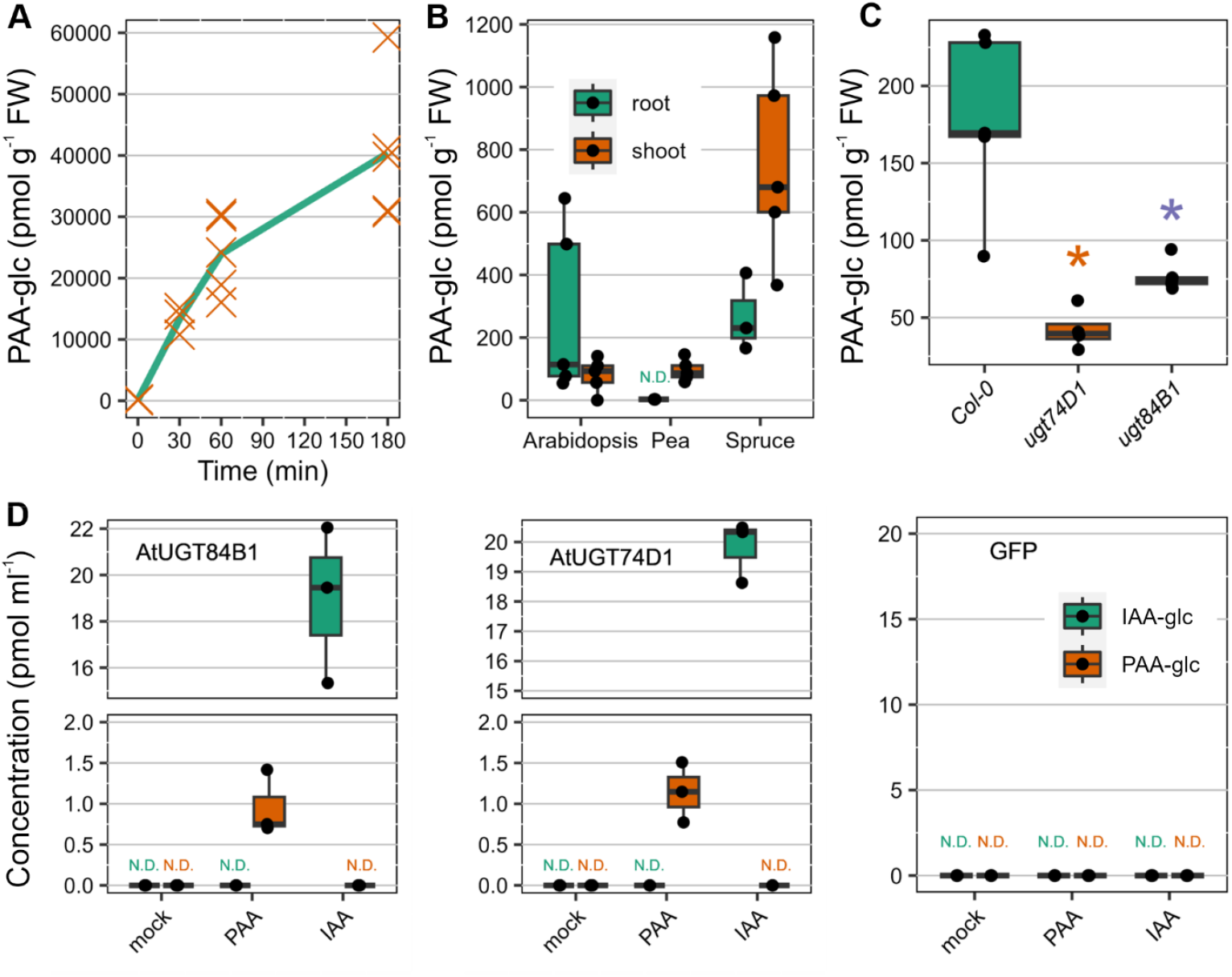
Presence and formation of PAA-glc in plants. Concentration (pmol g^-1^ FW) of PAA-glc after treatment of Arabidopsis with 20 μM PAA for 30, 60 and 180 min (A). Each orange cross represents an individual biological replicate (n=5). Endogenous levels of PAA-glc (pmol g^-1^ FW) in roots and shoots of Arabidopsis, pea and spruce (B). Levels of PAA-glc (pmol g^-1^ FW) in Col-0, ugt74d1 and ugt84b1 knockout Arabidopsis lines (C). Analysis of IAA-glc and PAA-glc synthesized by recombinant AtUGT84B1 and AtUGT47D1 produced by a bacterial assay (D). The cell lysate was incubated with 0.1 mM IAA or PAA and UGT cofactors for 5 h at 30°C. Cell lysate without treatment was used as a mock sample. The box plots show the upper and lower quartiles, with horizontal lines indicating medians, and each dot representing a single biological replicate. GFP-producing bacteria supplemented with PAA-AAs were used as controls. Statistically significant differences are indicated by asterisks, as determined by Student’s t-test (P ≤ 0.05). All plant profiling was performed in five biological replicates (n=5) and bacterial enzyme assays in three biological replicates (n=3). N.D., not detected.

The quantitative tissue-specific analysis revealed highest levels of PAA-glc in spruce shoots (almost 760 pmol g^-1^ FW). The levels around 270 pmol g^-1^ FW were determined in spruce and Arabidopsis roots. Pea and Arabidopsis shoots contained 95 and 80 pmol g^-1^ FW, respectively. In other tissues and species (maize, wheat, and *Physcomitrium patens*) PAA-glc was not detected (**Fig. 1B, Tab. 1**).

**Table 1:** PAA conjugate levels in various plant species. PAA conjugates were quantified (pmol g^-1^ FW ± SD; n=5) in roots, shoots and cotyledons of pea, wheat and maize, roots and shoots of Arabidopsis and spruce, and gametophores of Physcomitrium patens. <LOD, under the limit of detection.

| Species | Tissue | PAA | PAA-Asp | PAA-Glu | PAA-glc |
| --- | --- | --- | --- | --- | --- |
| Mean ± SD (pmol g<sup>-1</sup> FW) |  |  |  |  |  |
| *Arabidopsis thaliana* | shoot | 310.18 ± 23.25 | 581.03 ± 156.00 | 507.41 ± 130.00 | 80.21 ± 54.10 |
|  | root | 590.80 ± 273.62 | 1,896.87 ± 906.75 | 1,928.48 ± 868.86 | 277.82 ± 273.96 |
| Maize (*Zea mays*) | shoot | <LOD | 9.11 ± 2.56 | 10.18 ± 2.45 | <LOD |
|  | cotyledon | 422.34 ± 59.50 | 25.41 ± 21.87 | <LOD | <LOD |
|  | root | 185.05 ± 21.43 | 50.83 ± 7.29 | 99.54 ± 27.47 | <LOD |
| Wheat (*Triticum aestivum*) | shoot | 492.95 ± 98.83 | 16.91 ± 2.18 | <LOD | <LOD |
|  | cotyledon | 695.43 ± 504.86 | 432.38 ± 277.06 | 16.71 ± 11.46 | <LOD |
|  | root | 340.98 ± 112.34 | 523.48 ± 345.26 | <LOD | <LOD |
| Pea (*Pisum sativum*) | shoot | 1,113.51 ± 304.78 | 1,444.72 ± 466.83 | 54.06 ± 7.77 | 95.20 ± 34.25 |
|  | cotyledon | 310.19 ± 62.28 | 37,077.09 ± 12,823.16 | 1719.90 ± 413.77 | <LOD |
|  | root | 1,344.98 ± 169.59 | 1,095.36 ± 990.75 | 272.08 ± 53.63 | <LOD |
| Spruce (*Picea abies*) | shoot | 36.64 ± 13.17 | 1.62 ± 0.67 | 5.83 ± 0.34 | 755.73 ± 311.91 |
|  | root | 76.47 ± 18.58 | 1.94 ± 0.36 | 2.12 ± 0.24 | 267.79 ± 124.22 |
| *Physcomitrium patens* | - | 82.56 ± 36.08 | 1.82 ± 0.32 | 27.33 ± 3.46 | <LOD |

The glucosyltransferase UGT84B1 has been identified to be responsible for forming IAA-glc and PAA-glc *in vitro* (Aoi et al., 2020c). Similarly, UGT74D1 has been linked to the formation of oxIAA-glc (Tanaka et al., 2014; Brunoni et al., 2019; Mateo-Bonmatí et al., 2021), although its involvement in PAA metabolism has not been explored. To investigate whether these enzymes are involved in PAA-glc formation, we tested the conjugation activity of these enzymes by producing them in *E. coli* and using a bacterial assay designed to study various IAA catabolic enzymes (Brunoni et al., 2019; Brunoni et al., 2023a). Both UGT84B1 and UGT74D1 recombinant proteins showed the capability to produce IAA-glc and PAA-glc after exposure to 0.1 μM IAA and PAA, respectively (**Fig. 1D**). However, the activity of both glucosyltransferases towards PAA was only about 5% compared to IAA. To confirm their activity in plants, we explored PAA-glc content in Arabidopsis knockout lines *ugt84b1* and *ugt74d1*. Remarkably, we observed significantly lower levels of PAA-glc in both mutants compared to Col-0 (**Figure 1c**). In conclusion, both experiments demonstrated involvement of UGT84B1 and UGT74D1 in PAA glucosylation.

### Exploring novel PAA amide conjugates, their enzymatic synthesis and break down

Conjugates of IAA with various amino acids have been previously determined in plants. However, the only known amide conjugates of PAA in plants are those linked with Asp, Glu, and Trp (Sugawara et al., 2015; Staswick et al., 2017). In this study, in addition to identifying PAA-glc, we uncovered three previously unreported amide conjugates, PAA-Leu, PAA-Phe, and PAA-Val, in pea and wheat. The verification of newly identified endogenous conjugates relied on comparing their chromatographic retention times to those of synthetic standards (**Fig. S2)**. While endogenous steady-state levels of PAA-Leu, PAA-Phe, PAA-Trp, and PAA-Val in Arabidopsis were below the detection limits, feeding plants with 20 μM PAA promoted their *de novo* synthesis, resulting in detectable endogenous concentrations ranging from 0.5 to 1 pmol g^-1^ FW already after 30 min (**Fig. 2A**).

**Figure 2:**
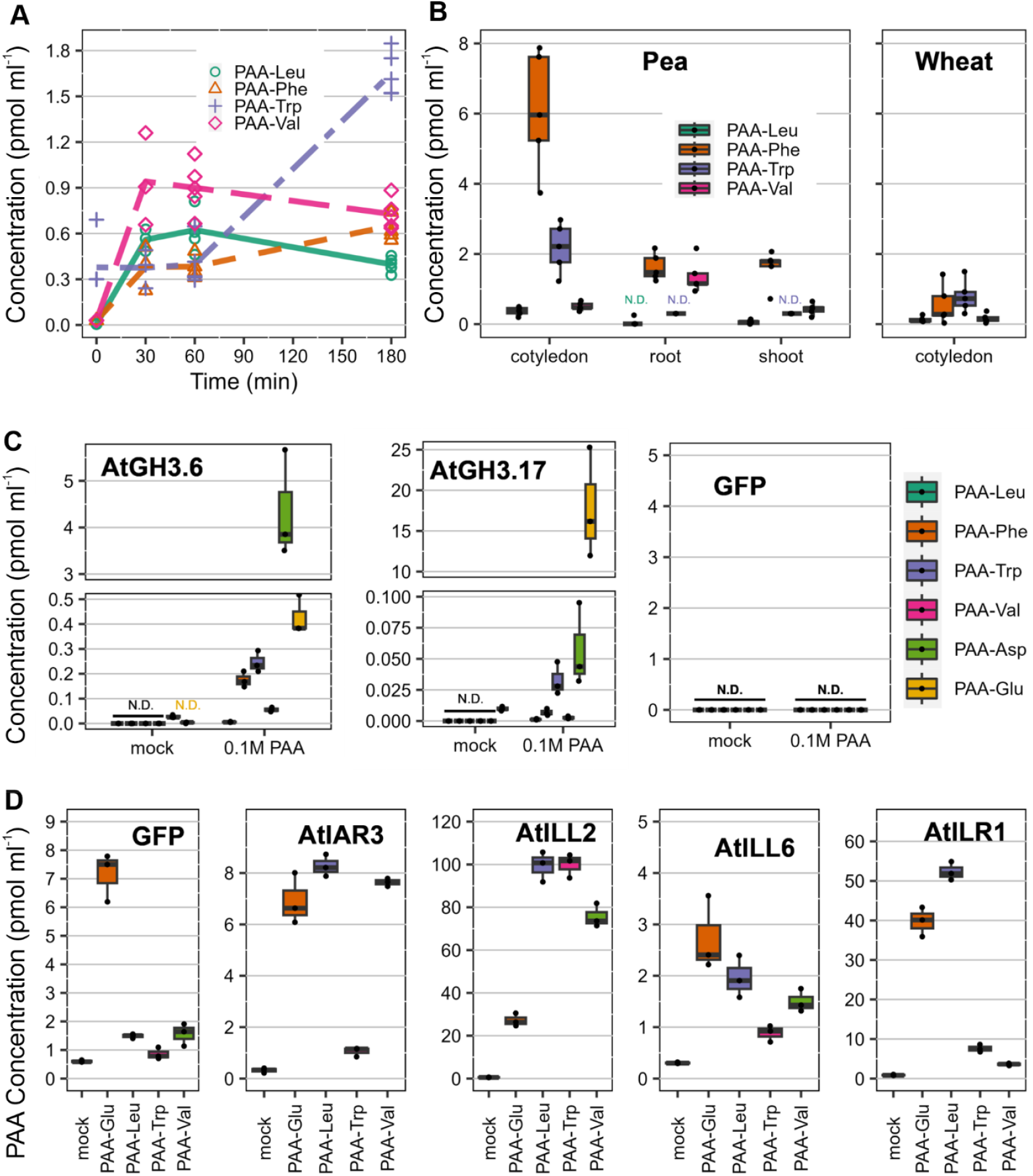
PAA amide conjugates abundance, formation and hydrolysis. The concentration levels (pmol g^-1^ FW) of PAA-Leu, PAA-Phe, PAA-Trp and PAA-Val were measured in Arabidopsis seedlings after treatment with 20 μM PAA for 30, 60 and 180 minutes (A). Each sign in a specific colour represents an individual biological replicate (n=5). The quantification of PAA-Leu, PAA-Phe, PAA-Trp and PAA-Val (pmol g^-1^ FW) was carried out in pea cotyledons, roots, and shoots, as well as in wheat cotyledons (B). The analysis of PAA-AAs synthesized by recombinant AtGH3.6 and AtGH3.17 in the bacterial assay. The cell lysate was incubated with 0.1 mM PAA and GH3 cofactors for 5 h at 30°C, and the formation of PAA-AAs was determined. As a control, cell lysate without PAA treatment was used (C). The hydrolysis of PAA-AAs to PAA was examined using a bacterial assay with recombinant AtIAR3, AtILL2, AtILL6 and AtILR2 (D). The lysate was incubated with 0.1 mM PAA-Leu, PAA-Trp, PAA-Val and PAA-Glu for 5 h at 30°C, and the levels of PAA were measured. A negative control was performed using GFP-producing bacteria, and a mock was performed using cell lysate without treatment. The box plots display medians as horizontal lines, upper and lower quartiles as boxes, and each dot represents a single biological replicate. All plant profiling was performed in five biological replicates (n=5) and bacterial enzyme assays in three biological replicates (n=3). N.D., not detected.

PAA-Leu, PAA-Phe, PAA-Val, and PAA-Trp were then quantified in different tissues of various plant species, with detectable concentrations observed only in pea and wheat (**Fig. 2B**). Pea cotyledons contained all four conjugates in concentrations ranging from 0.5 to 8 pmol g^-1^ FW. PAA-Phe, PAA-Val, and PAA-Leu were detected in roots, and PAA-Phe and PAA-Val were also found in shoots, all in concentrations below 2 pmol g^-1^ FW. All of them were identified in wheat cotyledon at concentrations not exceeding 2 pmol g^-1^ FW.

Previous works showed the involvement of IAA-conjugating GH3 enzymes in the metabolic regulation of PAA (Staswick et al., 2005; Westfall et al., 2016; Aoi et al., 2020b). To investigate the spectrum activity of GH3 enzymes to form PAA amino acid conjugates, recombinant AtGH3.6 and AtGH3.17 enzymes were tested with PAA using a bacterial assay (**Fig. 2C**). In the presence of 0.1 μM PAA, both enzymes formed all tested PAA amino acid conjugates. Nonetheless, AtGH3.6 predominantly conjugated PAA with Asp and AtGH3.17 with Glu.

Several IAA-AAs can be hydrolysed by members of the ILR1/ILL family, thus contributing to the free IAA pool in addition to *de novo* synthesis (LeClere et al., 2002; Rampey et al., 2004; Hayashi et al., 2021). To investigate whether PAA-AAs could undergo a similar level of regulation, we tested the possible hydrolysing activity of recombinant AtILL2, AtILL6, AtILR1, and AtIAR3 enzymes with 0.1 mM PAA-Leu, PAA-Val, PAA-Trp, or PAA-Glu and followed the formation of PAA (**Fig. 2D**). GFP-producing bacteria supplemented with PAA-AAs were used as controls. AtIAR3 preferentially hydrolysed PAA-Glu/-Leu/-Val, while AtILL6 did not exhibit any clear substrate preference. However, the amount of PAA formed by these enzymes was comparable to the GFP control, indicating the presence of endogenous substrate-associated machinery in *E. coli*, and thus, AtIAR3 and AtILL6 may not significantly contribute to the hydrolysis of these PAA amino acid conjugates. AtILL2 showed a pronounced preference for hydrolysing PAA-Leu/-Trp/-Val, while AtILR1 preferably hydrolysed PAA-Glu and PAA-Leu. Overall, our findings demonstrate the capability of ILR/ILL proteins to hydrolyse PAA amino acid conjugates.

### Crystal structures determination and docking of AAs with GH3s

Available crystal structures of AtGH3.6 with inhibitor (PDB 7VKA; Xie et al., 2022), AtGH3.5 with IAA (PDB 5KOD; Westfall et al., 2016), AtGH3.12 (PDB 4EPM and 4EQL, the latter containing salicylate; Westfall et al., 2012) or AtGH3.15 (PDB 6AVH; Sherp et al., 2018) contain AMP (as a product of ATP degradation) but not a free amino acid entering the ligase reaction with the second substrate (IAA/PAA). To understand the basis for amino acid specificity in GH3s, we solved the crystal structure of AtGH3.6 in the presence of AMP only, or AMP together with aspartate, up to 1.74 Å resolution. Data collection and refinement statistics are in **Tab. S4**. Superposition of both structures did not reveal rearrangement in the presence of amino acid (aspartate) ligand in the active site. The α-amino group of aspartate interacts with the phosphate group of AMP. The α-carboxyl group is H-bonded to the nitrogen atom of Leu175 and via two water molecules to the hydroxyl of Thr176 and main-chain oxygen of Phe158 (**Fig. 3A**). Moreover, both α-carboxyl and α-amino groups interact via a water molecule with Ser341 and the oxygen atom of Thr108 (not shown). The γ-carboxyl group is bound to Arg117, Lys160 and Ser455. Binding affinities for AtGH3.6 were analysed by MST. *K*_D_ values for Glu and Asp were 0.72 and 0.68 mM, respectively (**Fig. 3B**). Binding of Val and Leu was not measurable by MST. Affinities for IAA and PAA were similar and at around 60 μM concentration. The superposition of AtGH3.6 with AtGH3.5 shows the conservation of the amino-acid binding site and, thus, similar specificity (**Fig. 3C**).

**Figure 3:**
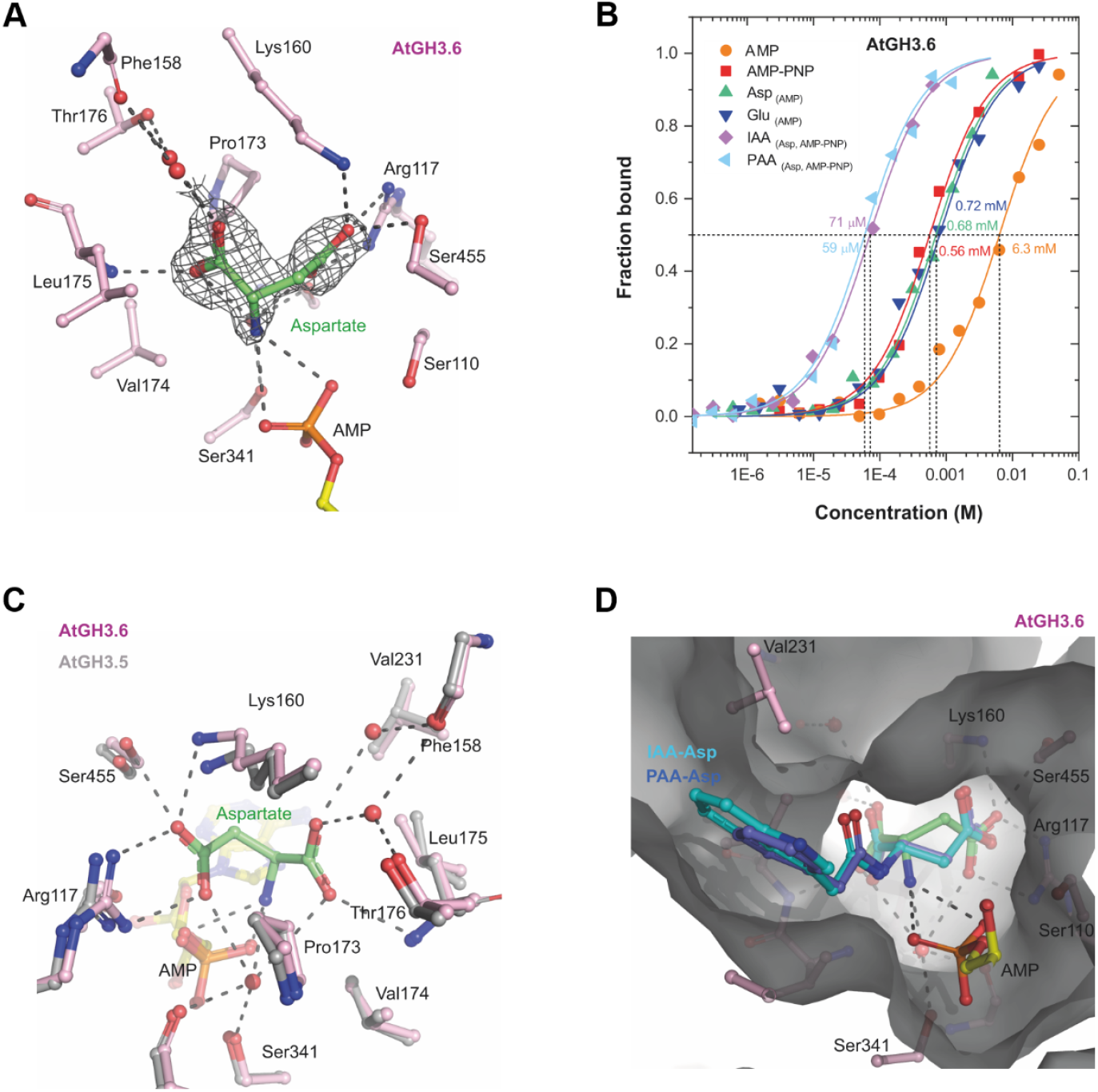
Binding interactions of aspartate in the active site of AtGH3.6. Binding of Asp (coloured in green) in AtGH3.6 (A). Surrounding residues are coloured pink and labeled. Ligand is depicted in its annealing Fo-Fc omit map (black mesh) contoured at 3.5 σ (PDB 9FXD). Binding affinity curves of AtGH3.6 for selected ligands, AMP, AMP-PNP, Glu, Asp, IAA-Asp and PAA-Asp (B). Data were measured by MST in 50 mM MES buffer pH 6.5, 1 mM MgCl_2_ and 0.05% Tween. Superposition of the amino acid binding site of AthGH3.6 (coloured in pink, PDB 9FXD) with AtGH3.5 structure (coloured in grey, PDB 5KOD) (C). Docked positions of IAA-Asp and PAA-Asp in the active site of AtGH3.6 (D). Docking calculation was performed by FLARE v 8.0.

Our attempts to obtain structures with bound products IAA-Asp or PAA-Asp were unsuccessful due to solubility issues of these ligands and a loss of crystal diffraction upon co-crystallization with AtGH3.6. Thus, we docked these two compounds as well as the amino acids in the active site of AtGH3.6 and AtGH3.5 by FLARE v 8.0 (Cheeseright et al., 2006; Bauer and Mackey, 2019; CRESSET, http://www.cresset-group.com/flare/) using the Lead Finder (LF) docking algorithm (**Tab. S5**). The LF ranking score and Gibbs free energy (ΔG) score for Asp and Glu were similar and mutually comparable, confirming their similar affinities observed by MST. The ΔG score estimates the free energy of protein-ligand binding for a given protein-ligand complex. Similar LF ranking scores were obtained for the IAA- and PAA-conjugates with Asp/Glu in line with similar affinities for IAA/PAA measured by MST. The best positions for IAA-Asp and PAA-Asp in the AtGH3.6 site are shown in **Fig. 3D**.

### PAA conjugates profiling in land plants

While previous studies have extensively examined PAA levels across various plant species and tissues (reviewed in Perez et al., 2023), information regarding its conjugate levels is mainly restricted to Arabidopsis (Sugawara et al., 2015; Aoi et al., 2020b). Therefore, we conducted a thorough tissue-specific profiling encompassing high-abundance PAA conjugates, namely PAA-Asp, PAA-Glu and PAA-glc, across diverse plant species of land plants (**Tab. 1**). Intriguingly, we observed significant variations in the PAA conjugate profile among species and even within tissues.

Remarkably, spruce and *Physcomitrium* exhibited lower levels of free PAA and PAA-Asp than other angiosperm representatives. Additionally, spruce represented the only species where the predominant conjugate was PAA-glc, accounting for around 95% of shoots’ PAA pool. Among other species, amide conjugates were more abundant. In moss, the level of PAA-Glu was more than 10 times higher than PAA-Asp, unlike in all other species. In both monocots, maize and wheat, cotyledons accumulated PAA the most. PAA-glc was not detected in these species. In *Arabidopsis*, most of the PAA conjugates were present in roots, primarily in the form of PAA-AAs. However, notable levels of free PAA and PAA-glc were also detected. In pea, PAA-Asp was the predominant storage form, with concentrations reaching approximately 40 nmol/g in cotyledons. Overall, pea stores a considerable amount of PAA in amide forms.

### Metabolic pathways of PAA display only partial functional redundancy

Previous studies have demonstrated the presence of functional redundancy within pathways of IAA metabolism (Porco et al., 2016; Mellor et al., 2016). Here, we aimed to investigate the dynamic changes in PAA and its conjugates in response to perturbations in specific metabolic pathways, providing insights into the mechanisms governing PAA homeostasis.

Arabidopsis *gh3sex* mutant seedlings, affected in the GH3-dependent PAA-AAs synthesis, were treated with 20 μM PAA for 30, 60, and 180 min, and the levels of PAA amide conjugates were compared with wild-type (Col-0). The PAA levels immediately increased after treatment, suggesting rapid uptake of exogenously applied PAA by plants (**Fig. 4A**). Subsequent analysis at 60 min post-treatment revealed significant differences in PAA concentrations between the *gh3sex* and Col-0 lines. Notably, while wild-type plants accumulated PAA-Asp over time, no PAA-Asp was detected in the mutant throughout the experiment (**Fig. 4B**), highlighting the dominant role of AtGH3.1-6 enzymes in the conjugation of PAA with Asp. The PAA-Glu levels were elevated in the *gh3sex* mutant at 60 and 180 min post-treatment (**Fig. 4C**). The increase in PAA-Glu levels in mutant could be attributed to compensatory mechanisms and functional redundancy of GH3 proteins (Porco et al., 2016; Brunoni et al., 2023a). Measurement of PAA-glc levels did not reveal any significant differences between wild-type and mutant (**Fig. 4D**).

**Figure 4:**
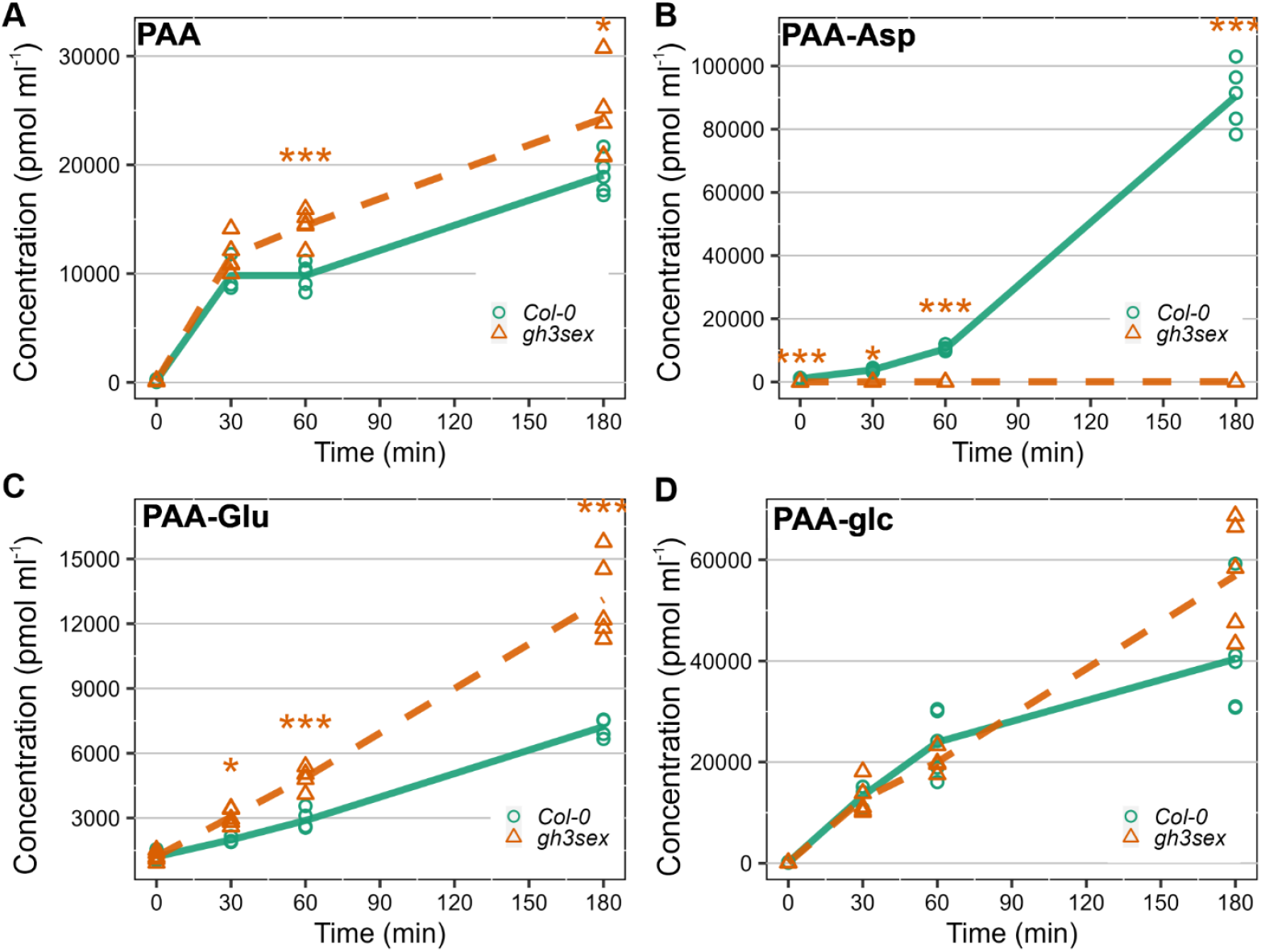
PAA metabolism in Arabidopsis gh3.1-6 knockout mutant. Arabidopsis line gh3.1,2,3,4,5,6 (gh3sex) and Columbia (Col-0) were treated with 20 μM PAA for 30, 60, and 180 min. The concentrations levels (pmol g^-1^ FW) of PAA (A), PAA-Asp (B), PAA-Glu (C), and PAA-glc (D) were measured in those four time points. Each sign in a specific colour represents an individual biological replicate (n=5). Asterisks indicate statistically significant differences between the Col-0 and mutant line in one time point, as determined by Student’s t-test (*, P ≤ 0.05; **, P ≤ 0.01; ***, P ≤ 0.001). The colour of the asterisk corresponds to the mutant line that is significantly different from Col-0.

To compare PAA metabolism across phylogenetically diverse plant species, we employed the synthetic GH3 inhibitor KKI, known for inhibiting the formation of IAA-AAs (Fukui et al., 2022). Anticipating a similar function in PAA metabolism, we investigated its effects in Arabidopsis, spruce and *Physcomitrium*, by treating them with 50 μM KKI, 5 μM PAA, or a combination of both.

In Arabidopsis, co-treatment with PAA and KKI resulted in reduced levels of PAA-Asp and PAA-Glu compared to PAA treatment alone, confirming the activity of KKI in inhibiting PAA-AAs formation. Interestingly, levels of free PAA remained unchanged between the two treatments (**Fig. 5A**). In spruce, the treatment with PAA confirmed predominant role of glucosylation in PAA metabolism. In agreement to Arabidopsis, KKI inhibited the GH3-mediated formation of both amide conjugates (**Fig. 5B**). Interestingly, in *Physcomitrium*, only formation of PAA-Glu was blocked by KKI in co-treatment with PAA, while PAA-Asp level was elevated (**Fig. 5C**). Notably, the formation of PAA-glc was not observed even after PAA treatment, suggesting that glucosylation does not occur in PAA metabolism in *Physcomitrium*.

**Figure 5:**
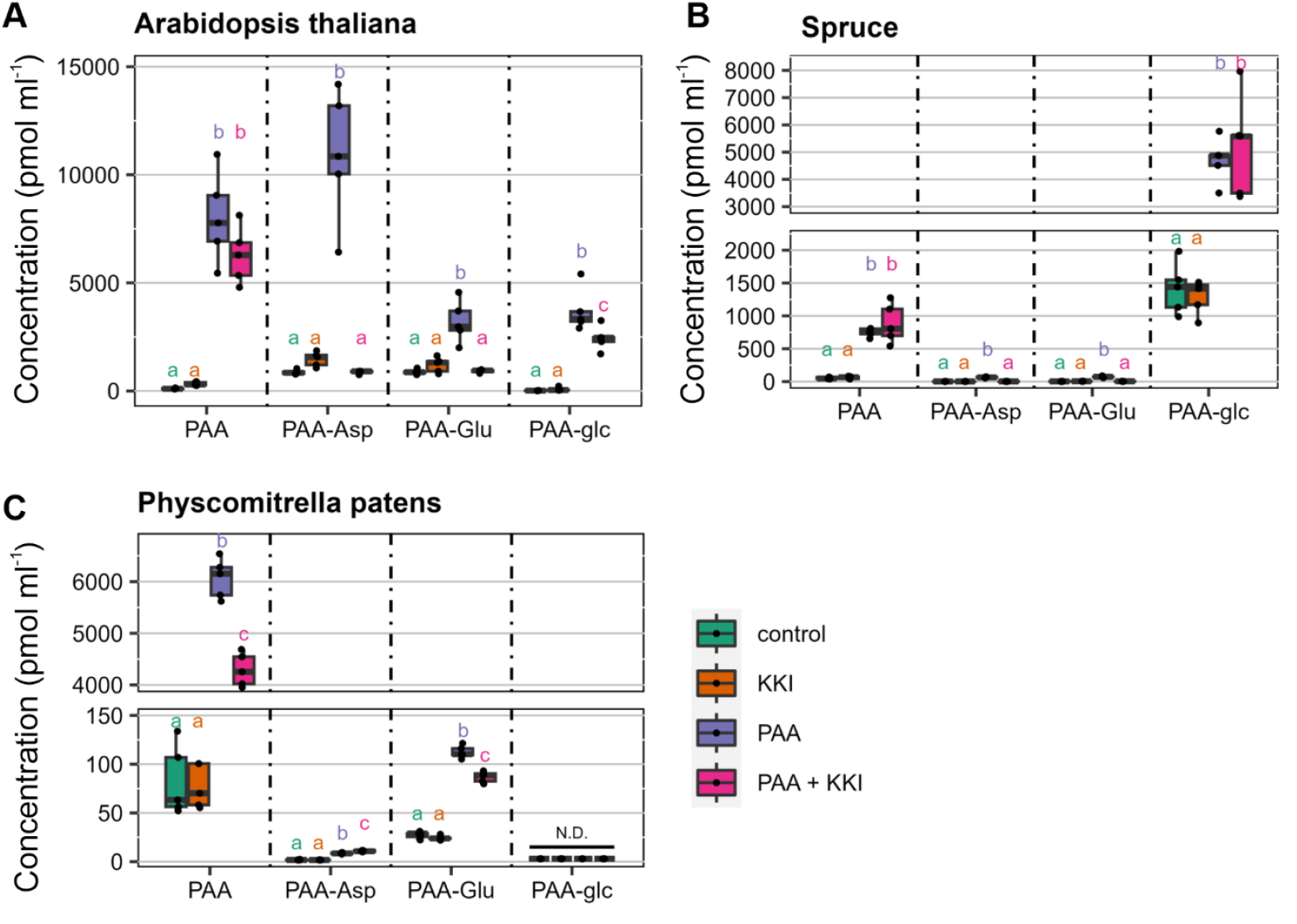
PAA metabolism after PAA and kakeimide (KKI) treatment in various plant species. Arabidopsis thaliana (A), spruce (B) and Physcomitrium p. (C) were treated with 50 μM KKI, 5 μM PAA or their combination for 1, 6 or 24 h respectively, with time depending on the species. The concentration (pmol g^-1^ FW) of PAA, PAA-Asp, PAA-Glu and PAA-glc was measured after the treatment. As a control, mock treated samples were used. The box plots display medians as horizontal lines, upper and lower quartiles as boxes, and each dot represents a single biological replicate (n=5). One-way ANOVA and Tukey’s post hoc test were applied to assess the differences between treatment groups. Different letters (a-c) indicate significant differences at the 5% level of significance (P ≤ 0.05). The colour of the letters corresponds with the colour of the boxplot. N.D., not detected.

## Discussion

Metabolism plays a pivotal role in maintaining auxin homeostasis, by ensuring optimal levels of biologically active hormone within the plant. While extensive research governing IAA metabolism has been done in previous years (Brunoni et al., 2020; Hayashi et al., 2021; Mateo-Bonmatí et al., 2021; Müller et al., 2021; Brunoni et al., 2023a; Hladík et al., 2023), the inactivation pathways of PAA remain largely uncharacterised. Thus far, only PAA-Asp, PAA-Glu, and PAA-Trp were identified in Arabidopsis (Sugawara et al., 2015; Staswick et al., 2017). However, there is no evidence about other conjugates or metabolic pathways, as oxidation of the phenyl ring is unlikely and the formation of glucosyl ester (PAA-glc) has only been demonstrated *in vitro* (Aoi et al., 2020c).

In our study we aimed to broaden the understanding of PAA metabolism by investigating novel conjugates and metabolic pathways (**Fig. 6**).

**Figure 6:**
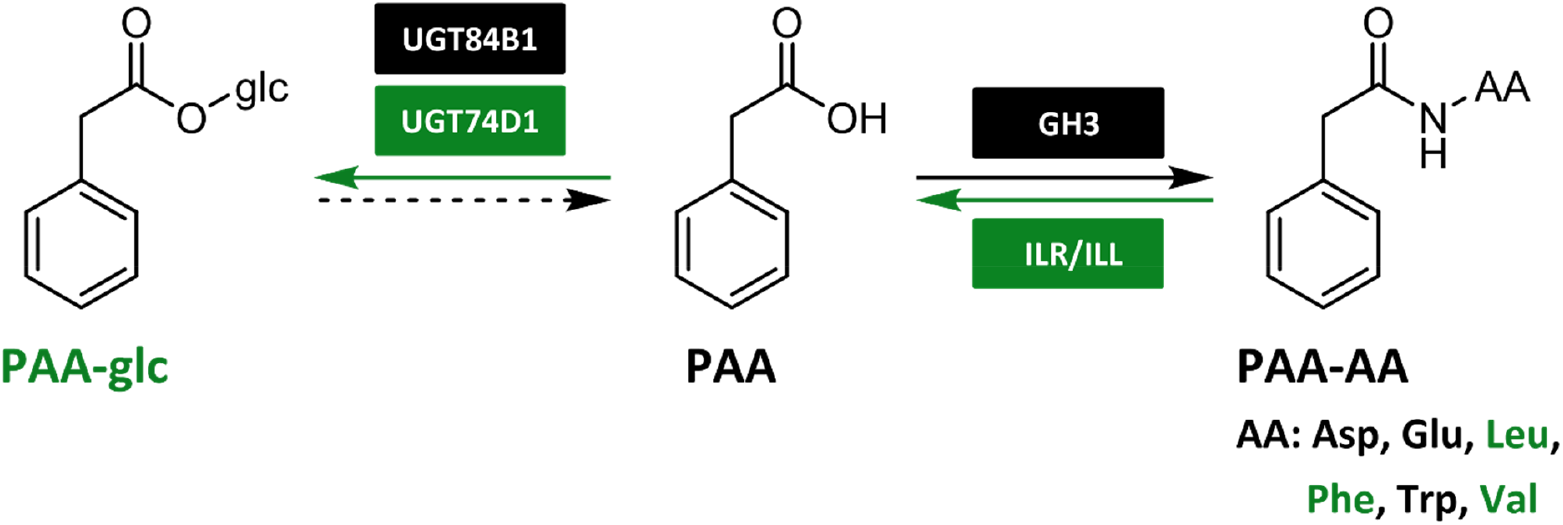
Updated scheme of PAA metabolism. Newly identified pathways and conjugates are highlighted in green. Dashed arrow represents putative metabolic step that is still not fully characterised. AA, amino acid; GH3, GRETCHEN HAGEN 3; ILR/ILL, IAA-LEUCINE RESISTANT 1/ILR1-LIKE proteins; PAA, phenylacetic acid; PAA-glc, PAA-glucose.

As a result of comprehensive multi-species screen, we confirmed occurrence of four novel PAA conjugates, PAA-glc, PAA-Leu, PAA-Phe and PAA-Val, in different plant species. The identity of endogenous conjugates was confirmed by comparison of their retention times with that of synthetic standards under the same chromatographic conditions. PAA-glc was found in Arabidopsis, pea and spruce, in concentrations ranging from 50 to 1000 pmol g^-1^ FW, with the highest levels observed in spruce shoots (**Fig. 1B**). However, even these high levels were close to the limit of detection of our method, likely due to poor ionisation of the molecule. It is plausible that PAA-glc may also be present in other studied species, however below the limit of detection. Presence of newly identified amide conjugates, PAA-Leu, PAA-Phe, and PAA-Val, was observed only in pea and wheat in low concentrations ranging from 0.5 to 8 pmol g^-1^ FW (**Fig. 2B**). These findings align with the low levels of IAA and oxIAA conjugates with amino acids other than Asp and Glu quantified previously in various plants (Kowalczyk & Sandberg, 2001; Pěnčík et al., 2009; Hladík et al., 2023). While Staswick et al., (2017) observed high levels of PAA-Trp (approximately 30 pmol g^-1^ FW) in Arabidopsis tissue, under our experimental conditions PAA-Trp was not detected in Arabidopsis, being only determined in pea and wheat cotyledons. Although PAA conjugates with Leu, Phe, Trp, and Val were not found in Arabidopsis, the capability of Arabidopsis GH3 proteins to catalyse their synthesis was proved by a bacterial enzymatic assay (**Fig. 2C**). To further validate the formation of newly identified metabolites *in planta*, we conducted feeding experiments by supplying exogenous PAA to Arabidopsis seedlings. This led to rapid synthesis of PAA-glc (**Fig. 1A**) as well as all three novel amide conjugates (**Fig. 2A**), underscoring the role of these conjugates in maintaining PAA homeostasis.

Formation of PAA-glc by enzyme UGT84B1 has already been shown *in vitro* (Aoi at el., 2020b). However, other glucosyltransferases can also be involved in formation of IAA/oxIAA-glc, such as UGT74D1 (Jackson et al., 2001; Brunoni et al., 2019; Mateo-Bonmatí et al., 2021). Thus, we tested the conjugation activity of this protein in bacterial assay designed to study IAA enzymatic inactivation (Brunoni et al., 2019; Brunoni et al., 2023a) and proved the capability of AtUGT74D1 to produce PAA-glc (**Fig. 1D**). Additionally, we quantified PAA-glc in *ugt84b1* and *ugt74d1* knockouts and demonstrated the involvement of both proteins in formation of PAA-glc *in planta* (**Fig. 1C**).

The GH3-mediated formation of IAA amide conjugates is a well described mechanism (Staswick et al., 2005; Zhang et al., 2018). The role of GH3s in PAA metabolism was also indicated for PAA-Asp and PAA-Glu formation (Sugawara et al., 2015; Westfall et al., 2016; Aoi et al., 2020b). To investigate the role of GH3s in formation of other PAA-AAs, we performed bacterial enzyme assays with AtGH3.6 and AtGH3.17 (**Fig. 2C**). Results indicated that both GH3 proteins are capable of synthesizing all tested PAA conjugates, with AtGH3.6 displaying a preference for Asp and AtGH3.17 for Glu as substrates, aligning with previous assays with IAA conjugation (Brunoni et al., 2019; Brunoni et al., 2023a). Accordingly, the crystal structure of AtGH3.6 in the presence of Asp (PDB 9FXD) indicates hydrogen bonding of the Asp side chain with three protein residues (Arg117, Lys160, Ser455), favouring Asp binding (**Fig. 3**). Of note, superposition of 9FXD structure with that of AtGH3.5 with bound AMP and IAA (PDB 5KOD) shows that these three conserved residues are positioned similarly, favouring Asp binding. Final activity and thus specificity depends not only on affinities but also on transition state stability and association and dissociation rate of ES complex plus availability of particular ligands, which can lead to differences among GH3 family enzymes among various species. Furthermore, we investigated the putative hydrolysis of PAA-AAs by ILR/ILL amidohydrolases (**Fig. 2D**), as previously described for IAA conjugates (Bartel & Fink, 1995; Davies et al., 1999; LeClere et al., 2002; Hayashi et al., 2021). Bacterial enzyme assays with AtILL2, AtILL6, AtILR1, and AtIAR3 revealed that PAA-AAs can be hydrolysed into free PAA, indicating storage function of PAA amino acid conjugates.

To elucidate evolutionary aspects of PAA metabolism, we conducted profiling of PAA and its major conjugates, PAA-Asp, PAA-Glu, and PAA-glc across a spectrum of phylogenetically diverse land plants. Our study encompassed representatives such as the moss *Physcomitrium*, spruce as a representative of gymnosperm trees, dicots represented by Arabidopsis and pea, and two monocots, maize and wheat (**Tab. 1**).

According to our findings, PAA levels largely align with previous studies, revealing consistent PAA levels in Arabidopsis and pea tissues (Wightman & Lighty, 1982; Sugawara et al., 2015). Notably, comparison with our previous IAA quantifications (Hladík et al., 2023) as well as with earlier reports indicates significantly higher PAA levels in most plant species and tissues. PAA conjugate profiling revealed PAA-AAs as major metabolites across all studied plants except spruce, where PAA-glc concentrations were notably higher compared to PAA amides. This finding, together with results obtained from PAA feeding experiment (**Fig. 5B**) and previously reported evidence that IAA glucosylation is the main pathway to maintain IAA homeostasis (Brunoni et al., 2020), suggests that glucosylation serves as the preferred pathway for PAA and IAA inactivation in spruce. Remarkably, the exceptionally high PAA-AAs concentrations in pea mirror elevated levels of IAA-AAs and oxIAA-AAs in pea tissues (Hladík et al., 2023), suggesting analogous metabolic regulation of both auxins. Although PAA conjugation pathways share similarities with those of IAA, the oxidation to oxIAA that serves as a degradation mechanism of IAA and IAA-AAs (Hayashi et al., 2021), represents notable difference between IAA and PAA metabolism. Both oxIAA and its glucosyl ester are considerably more abundant among IAA metabolites (Kai et al., 2007a; Pěnčík et al., 2018; Hladík et al., 2023). In contrast, PAA-Asp and PAA-Glu exhibit substantially higher accumulation compared to IAA amide conjugates and represent predominant PAA metabolites in Arabidopsis (**Tab. 1**).

To investigate putative functional redundancy in PAA inactivation between GH3s and UGTs, we explored PAA metabolism using GH3-deficient Arabidopsis mutant and the synthetic GH3 inhibitor KKI (Fukui et al., 2022). Following the application of PAA, the synthesis of PAA-Asp was dramatically reduced in *gh3sex* (**Fig. 4**). This reduction was partially compensated by an increased conjugation of PAA to Glu. Notably, the deficiency in GH3-mediated conjugation was not compensated by glucosylation, mirroring observations seen with IAA (Porco et al., 2016). This observation was further confirmed by the co-treatment of Arabidopsis, spruce and *Physcomitrium* with PAA and KKI, where no metabolic compensation between GH3s and UGTs was observed (**Fig. 5**).

In conclusion, our investigation of PAA metabolism has provided valuable insights into the metabolic pathways governing PAA homeostasis in land plants. It appears that there may be other metabolic pathways of PAA that have yet to be discovered, as many have been found in bacteria (Schneider et al., 1997; Navarro-Llorens et al., 2005; Teufel et al., 2010). However, through the identification of novel PAA conjugates and the elucidation of metabolic pathways, we have expanded our understanding of the mechanisms maintaining PAA homeostasis and demonstrated the complexity and species-specific nature of PAA metabolism.

## Materials and Methods

### Reagents and standards

Plant agar and Murashige & Skoog media were purchased from Duchefa (Haarlem, Netherlands). Hypergrade purity methanol for HPLC-MS/MS analysis and all other chemicals were purchased from Lach-Ner (Neratovice, Czech Republic), Merck KGaA (Darmstadt, Germany), and Sigma-Aldrich (St. Louis, MA, USA). Standards for PAA and ^13^C_6_-labeled PAA were purchased from Merck KGaA (Darmstadt, Germany). IAA-glc and [^13^C_6_]IAA-glc were synthesised according to (Kai et al., 2007a; Kai et al., 2007b) with minor modifications. Selected L-amino acid (Val, Leu, Phe, Trp, Asp, Glu) conjugates of phenylacetic acid (PAA), including isotopically labelled standards [^13^C_6_]PAA-Asp and [^13^C_6_]PAA-Glu, were prepared according to Magnus et al., (1997). PAA-glc was synthesized adopting reaction conditions from Takeuchi et al., (2020). Detailed procedures for the synthesis of PAA conjugates are described in Supplementary Information (**Methods S1**).

### Plant material and growth conditions

*Arabidopsis thaliana* seeds ecotype Columbia 0 (Col-0) were used as wild-type for all the experiments. Knockout mutant lines *gh3*.*1,2,3,4,5,6* (*gh3sex*) (Porco et al., 2016), *ugt74d1* and *ugt84b1* (Mateo-Bonmatí et al., 2021) were obtained from prof. Karin Ljung (UmeÅ Plant Science Centre, Sweden). *Arabidopsis thaliana* (L.), maize (*Zea mays* L.), pea (*Pisum sativum arvense* L.), and wheat (*Triticum aestivum* L.) were cultivated as previously published in Hladík et al., (2023). Gametophores from *Physcomitrium patens* and spruce (*Picea abies* L. Karst) plants were cultivated as described in Brunoni et al., (2023a). All the plants were harvested (≈ 10 mg/FW; *Fresh Weight*) at growth stage 1.0 according to the Biologische Bundesanstalt, Bundessortenamt und Chemische Industrie (BBCH) scale (Tottman, 1987; Lancashire et al., 1991; Boyes et al., 2001) (except *Physcomitrium*, which was harvested three weeks after the last gametophores transfer to fresh medium).

### Feeding experiments

For PAA treatments, seven days after germination (7 DAG) Arabidopsis seedlings (Col-0 and *gh3sex*) grown under the same conditions as described above were harvested, washed in ultrapure water, and transferred to liquid medium (½ MS medium, 1% sucrose, pH 5.7) supplemented with 20 μM PAA. Plants were shaken gently in the dark at 22°C and harvested after 0.5, 1, and 3 h. For kakeimide (KKI) (Fukui et al., 2022) treatments, 7 DAG Arabidopsis seedlings, 14 DAG spruce plants, and 3-week-old *Physcomitrium* gametophores were transferred to sterile liquid Knop medium for 1, 6, and 24 h, depending on the species, according to Fukui et al., (2022) and Brunoni et al., (2023a) and then supplemented with 5 μM PAA, 50 μM KKI or a combination of 5 μM PAA with 50 μM KKI. Mock-treated Arabidopsis, moss and spruce plants were used as controls. Plants were harvested in five biological replicates per time point (≈ 10 mg/FW), immediately snap-frozen in liquid nitrogen, and stored at -80°C.

### Cloning, protein production, and bacterial enzyme assay

*Escherichia coli* BL21 (DE3) strains expressing recombinant AtGH3s, AtUGTs, and AtILR1/ILLs used in this work were previously generated (Brunoni et al., 2019; 2023b). Recombinant protein production and enzymatic assay of AtGH3s, AtUGTs, and AtILR1/ILLs were performed as described previously by Brunoni et al., (2019; 2023a; 2023b). For the amino acid conjugation assay, 500 μl of clarified cell lysate from AtGH3.6- or AtGH3.17-producing bacterial cultures was incubated with GH3 cofactors and with or without 0.1 mM PAA. For the glucose conjugation assay, 500 μl of clarified cell lysate from AtUGT84B1- or AtUGT74D1-producing bacterial cultures was incubated with UGT cofactors and with or without 0.1 mM PAA/IAA. For the hydrolysis assay, 500 μl of clarified cell lysate from AtILL2-, AtILL6-, AtILR1-, or AtIAR3-producing bacterial cultures was incubated with 1 mM MgCl_2_ and with or without 0.1 mM PAA-Leu, PAA-Trp, PAA-Val or PAA-Glu. GFP-producing bacterial cultures were used as negative controls. The enzymatic activity of the recombinant proteins was tested for 5 h at 30°C with constant shaking at 50 rpm in darkness and repeated in three biological replicates.

### PAA conjugate profiling

Extraction and purification of PAA conjugates followed the methodology described by Hladík et al., (2023) with modifications. Samples containing ≈ 10 mg of fresh weight tissue were extracted in 1 ml of an ice-cold sodium phosphate buffer supplemented with 0.1% diethyldithiocarbamic acid sodium salt. A mixture of isotopically labelled internal standards (IS) was added to the samples, including [^13^C_6_]PAA (10 pmol), [^13^C_6_]PAA-Asp (5 pmol), [^13^C_6_]PAA-Glu (5 pmol). The samples were homogenised using an MM400 bead mill (Retsch GmbH, Haan, Germany) with three zirconium oxide beads. The samples were then incubated on a rotary shaker (15 min, 27 rpm, 4°C) and then centrifuged (10 min, 206 642 *g*, 4°C). From the supernatant, 200 μl were acidified with 1M HCl to pH 2.7 and subjected to purification by in-tip micro solid-phase extraction (in-tip μSPE) utilizing a combination of HLB AttractSPE™ (Affinisep, Le Houlme, France) and SDB-XC Empore™ (3M, MN, USA) sorbents. The multi-StageTip microcolumns were activated sequentially with 50 μl of acetone (centrifugation 10 min, 3 846 *g*, 8°C), 50 μl of methanol (10 min, 3 846 *g*, 8°C), and 50 μl of water (15 min, 4 654 g, 8°C). The acidified samples were then applied to the activated microcolumns (30 min, 16 961 *g*, 8°C), washed with 50 μl of 0.1% acetic acid (20 min, 9 846 *g*, 8°C), and eluted with 50 μl of 80% methanol (20 min, 8 653 *g*, 8°C). After elution, samples were evaporated to dryness under vacuum and stored at -20°C until HPLC-MS/MS analysis.

Evaporated samples were reconstituted in 30 μl of 10% methanol prior to analysis on an HPLC-MS/MS system consisting of a 1260 Infinity LC II system (Agilent Technologies, Santa Clara, CA, USA) equipped with a reversed-phase chromatographic column (Kinetex C18 100 Å, 50 × 2.1 mm, 1.7 μm; Phenomenex, CA, USA) and coupled to a 6495B Triple Quadrupole LC/MS system (Agilent Technologies, CA, USA). The mobile phase consisted of deionised water (A) and methanol (B) supplemented with 0.1% acetic acid. The chromatographic analysis was carried out for 18 min at a flow rate of 0.3 ml/min. The elution of auxin metabolites was achieved using a gradient: 0 min - 10% B, 11.5 min - 60% B, 11.75 min - 99% B, 14.75 min - 99% B, 15 min - 10% B. During analysis, samples were stored in an autosampler at 4°C, with the column maintained at 40°C, and 10 μl of each sample was injected.

Individual analytes were detected using the MS instrument operating in negative electrospray ionisation (ESI-) mode with optimised parameters: nebuliser pressure at 25 psi, drying gas flow rate and temperature set at 14 l min^-1^ and 130°C respectively, sheath gas flow rate and temperature set at 12 l min^-1^ and 400°C respectively, capillary voltage set at 3.0 kV, and nozzle voltage maintained at 0 V. The measured analytes were detected and quantified by diagnostic multiple reaction monitoring (MRM) transitions of precursor and appropriate product ions using optimal collision energies and a dwell time of 50 ms, as described in **Tab. S1**. Raw data analysis was performed using Mass Hunter software (Agilent Technologies, CA, USA).

For method validation, 15-point calibration curve was prepared ranging from 9 amol to 90 pmol and the limit of detection (LOD) (S/N ratio > 3) was calculated as well as the dynamic linear range (**Tab. S1**). Validation protocol followed approach published in Hladík et al., (2023). Arabidopsis and pea plants were harvested at the growth stage 1.0 and spiked by 0, 1, and 10 pmol of authentic PAA standards (0, 10, and 50 pmol for PAA-glc), and 5 pmol of IS ([^13^C_6_]PAA, [^13^C_6_]PAA-Asp, [^13^C_6_]PAA-Glu). All samples were then extracted and purified as described above and measured by HPLC-MS/MS. After the measurement, analytes accuracy (percentage bias) and precision (relative standard deviation in %) was calculated (**Tab. S2**; **Tab. S3**).

### Statistical analyses

All analyses were performed using R statistical software (version 4.3.2; R Core Team, 2021) within the RStudio environment (version 2023.12.0.369; Posit team, 2023). The following packages were used for statistical analysis and graph generation: dplyr (Wickham et al., 2023a), ggplot2 (Wickham, 2016), ggbreak (Xu et al., 2021), multcomp (Hothorn et al., 2008), multcompview (Graves et al., 2019), and readxl (Wickham & Bryan, 2023b).

One-way ANOVA was used to assess differences between control and experimental variants. Significant differences detected at the 95% confidence level were subjected to Tukey’s *post-hoc* test. Values under the limit of detection (LOD) were replaced with 0.66-fold of the respective LOD value. For data visualisation, box-and-whisker plots were generated showing the median (centre line), upper and lower quartiles (box limits) and maximum and minimum values, with individual dots representing each biological replicate.

### Expression and purification of AtGH3.6 protein

A protein expression construct harbouring the coding sequence of AtGH3.6 (TAIR accession no. AT5G54510) was previously generated (pETM11-AtGH3.6, Brunoni et al., 2019), transformed into E. coli Rosetta (DE3) cells, and grown in Luria broth containing 50 μg/ml kanamycin and 20 μg/ml chloramphenicol to OD_600nm_ ∼0.8. Protein expression was induced with 1 mM isopropyl-b-D-thiogalactopyranoside (IPTG) overnight at 20°C. Cell lysis and protein extraction were performed as described in Brunoni et al. (2023a). The recombinant protein was purified on a Nickel-HiTrap IMACFF column (Cytiva Life Sciences, Marlborough, MA, USA) on an NGC Medium-Pressure Liquid Chromatography System into 50 mM HEPES (pH 7.5) and 300 mM NaCl. The His-tagged protein was further purified by anion exhange chromatography on a Resource Q column (Cytiva Life Sciences, Marlborough, MA, USA). Equilibration buffer was as follows: 50 mM HEPES (pH 7.5) and 1 mM MgCl_2_. The elution buffer additionally contained 1 M NaCl. The purified protein was desalted by diafiltration using 50-kDa-cutoff Centricon filters (Millipore, Bedford, MA). Protein concentration was determined by Bradford assay with bovine serum albumine (BSA) as the standard.

### Crystallization of AtGH3.6 and structure determination

Crystallization conditions for AtGH3.6 were screened using Qiagen PEGS II suite kit (www.qiagen.com). Crystals of AtGH3.6 were obtained in hanging drops by mixing equal volumes of i) enzyme solution (7.1 mg ml^-1^ for crystal with AMP and Asp, 8,2 mg ml^-1^ for crystal with AMP, in 20 mM HEPES buffer pH 7.5, 100 mM NaCl, 1 mM MgCl_2_, 1% glycerol) containing either 10 mM AMP or 10 mM AMP with 1 mM sodium aspartate, and ii) a precipitant solution containing 100 mM MES pH 6.5, 0.6 M NaCl and 18% (for crystal with AMP and Asp) or 20% (crystal with AMP) PEG 4000. Crystals were transferred to a cryoprotectant solution composed of the mother liquor supplemented with 20% PEG 400 and flash-frozen in liquid nitrogen. Diffraction data were collected at 100 K on the PROXIMA 1 and 2 beamlines at the SOLEIL synchrotron (www.synchrotron-soleil.fr). Intensities were integrated using the XDS program (Kabsch, 2010) and further reprocessed by Staraniso (Tickle et al., 2016). Data quality was assessed using the correlation coefficient *CC*_1/2_ (Karplus and Diederichs, 2012) (**Tab. S4**). Crystal structures were determined by performing molecular replacement with Phaser (McCoy et al., 2007) using the structure of AtGH3.5 (PDB 5KOD, Westfall et al., 2016) as a search model. Models were refined with NCS restraints and TLS using Buster 2.10 (Bricogne et al., 2011) and with ligand occupancies set to 1. Electron density maps were evaluated using COOT (Emsley and Cowtan, 2004). MolProbity was used for structure validation (Chen et al., 2010). Molecular graphics images were generated using PYMOL v 3.0 (www.pymol.org). Ligand interactions were analysed using Discovery Studio Visualizer (BIOVIA, San Diego, USA).

### Docking of Asp/Glu conjugates into the active site of AtGH3.6 and AtGH3.5

*In silico* docking was performed to compare the binding of Asp/Glu and their IAA and PAA conjugates into the active site of AtGH3.6 (PDB 9FXD, this work) and AtGH3.5 (PDB 5KOD, Westfall et al., 2016) by FLARE v 8.0 (Cheeseright et al., 2006; Bauer and Mackey, 2019; CRESSET, http://www.cresset-group.com/flare/). The proteins were prepared for docking using rule-based protonation predicted for pH 7.0 and intelligent capping. Energy grids for docking were 20 × 20 × 20 Å in dimension and centered on the amino group of co-crystallized aspartate ligand in the structure AtGH3.6. Docking calculations were carried out by the Lead Finder docking algorithm, with three independent docking runs and keeping the best poses overall (Kuhn et al., 2020). The resulting ligand orientations and conformations were scored based on their binding free energies and the Lead Finder rank score (Stroganov et al., 2008).

### Microscale thermophoresis (MST) affinity measurements

The MST method was used to determine the binding affinity of various amino acids, AMP-PNP (adenylyl-imidodiphosphate) and AMP as well as IAA and PAA ligands. Proteins were fluorescently labeled with RED-tris-NTA dye (www.nanotemper-technologies.com) using a 1:1 dye/protein molar ratio. The labeled protein was adjusted to 100 nM in 50 mM MES buffer pH 6.5, 1 mM MgCl_2_ and 0.05% Tween. Measurements were performed in premium capillaries on a Monolith NT.115 instrument at 30°C with 5 sec/30 sec/5 sec laser off/on/off times (medium MST power), respectively, with continuous sample fluorescence recording.

## Data availability

The atomic coordinates and structure factors have been deposited in the Protein Data Bank (www.wwpdb.org) under accession codes 9FWD for AtGH3.6 with AMP and 9FXD for AtGH3.6 with AMP and aspartate. The data that support the findings of this study are openly available in Zenodo at https://doi.org/10.5281/zenodo.13587370.

## Author contributions

ON and AP conceived the project. PH and AP performed method development and optimization. PH and FB grew and sampled the plants. PH and AP conducted the purification and quantification of auxin metabolites. FB performed bacterial enzymatic assays. AZ and MZ synthesized all new PAA conjugates standards. PH, ON, DK, and AP analysed and interpreted the data. JB, FB, DK, NF, and PB performed crystallization and docking experiments. PH prepared the manuscript draft. PH and AP wrote the article with input from all authors. All authors contributed to the article and approved the submitted version.

