## Supplemental information for "Phenylacetic acid metabolism in land plants: novel pathways and metabolites"

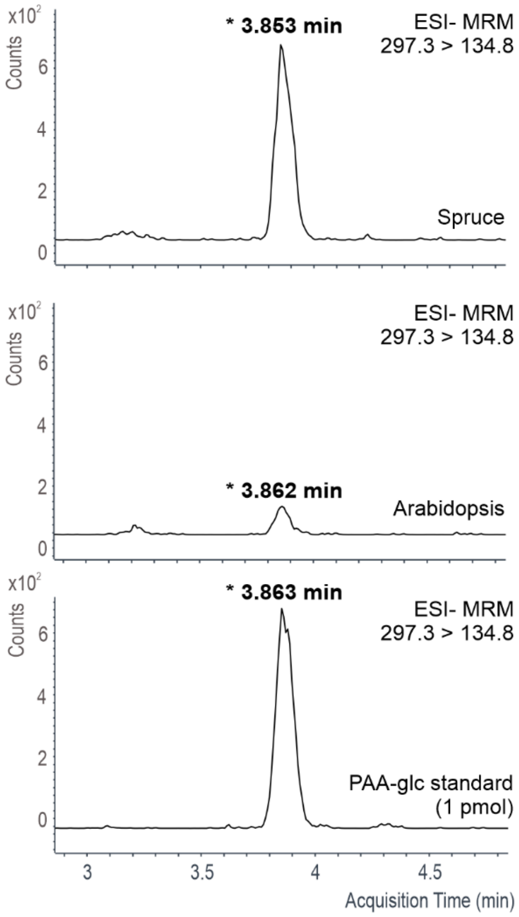


**Figure S1: Representative MRM chromatograms of PAA-glc.** Retention times of PAA-glc in spruce and Arabidopsis extracts containing 2 mg FW of the tissue and 1 pmol of synthetic reference standard.

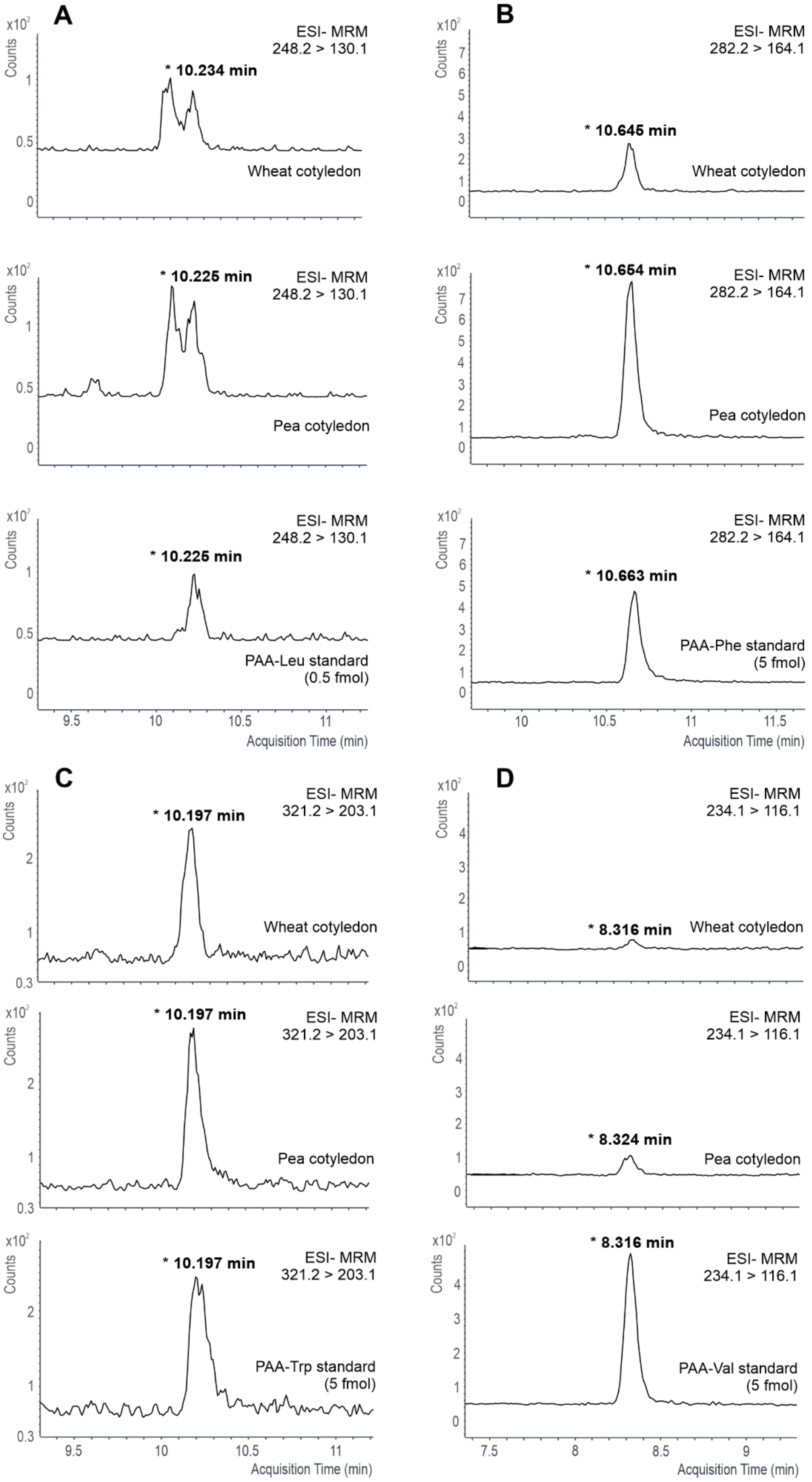


**Figure S2: Representative MRM chromatograms of PAA-AAs.** Retention times of PAA-Leu (A), PAA-Phe (B), PAA-Trp (C) and PAA-Val (D) in wheat and pea cotyledons extracts containing 2 mg FW of the tissue and 1 pmol of synthetic reference standard.

**Table S1: Conditions and parameters of HPLC-MS/MS method.** For each PAA metabolite and its corresponding internal standard (IS) diagnostic MRM transition and collision energies (CE) were optimized. Additionally, retention time (RT), limit of detection (LOD), linear range and coefficient of determination (R2) were measured and calculated. Analytes were detected by the MS instrument with optimised conditions as described: nebulizer pressure, 25 psi; drying gas flow and temperature, 14 l $min^{-1}$ and 130°C; sheath gas flow and temperature, 12 l $min^{-1}$ and 400°C; capillary voltage, 3.0 kv; nozzle voltage, 0 V.

| Compound | MRM transition | IS | MRM transition | CE (V) | retention time (min) | LOD (pmol) | linear range (pmol) | $R^2$ |
|---|---|---|---|---|---|---|---|---|
| **PAA** | 135.1 > 91.0 | [$^{13}C_6$]PAA | 141.1 > 97 | 2 | 5 | 4.5 $10^{-2}$ | 4.5 $10^{-2}$ - 90 | 0.9965 |
| **PAA-Asp** | 250.1 > 132.0 | [$^{13}C_6$]PAA-Asp | 256.2 > 132 | 10 | 2.9 | 4.5 $10^{-3}$ | 4.5 $10^{-3}$ - 90 | 0.9973 |
| **PAA-glc** | 297.3 > 91.0 | [$^{13}C_6$]PAA-Glu | 270.2 > 146 | 19 | 3.9 | 4.5 $10^{-2}$ | 9.0 $10^{-2}$ - 90 | 0.9904 |
| **PAA-Glu** | 264.1> 146.0 | [$^{13}C_6$]PAA-Glu | 270.2 > 146 | 12 | 3.7 | 9.0 $10^{-2}$ | 9.0 $10^{-2}$ - 90 | 0.996 |
| **PAA-Leu** | 248.2 > 130.1 | [$^{13}C_6$]PAA-Glu | 270.2 > 146 | 12 | 10.2 | 9.0 $10^{-5}$ | 9.0 $10^{-5}$ - 9 | 0.9985 |
| **PAA-Phe** | 282.2 > 164.1 | [$^{13}C_6$]PAA-Glu | 270.2 > 146 | 14 | 10.7 | 4.5 $10^{-4}$ | 4.5 $10^{-4}$ - 9 | 0.998 |
| **PAA-Trp** | 321.2 > 203.1 | [$^{13}C_6$]PAA-Glu | 270.2 > 146 | 14 | 10.2 | 4.5 $10^{-3}$ | 4.5 $10^{-3}$ - 9 | 0.9974 |
| **PAA-Val** | 234.1 > 116.1 | [$^{13}C_6$]PAA-Glu | 270.2 > 146 | 12 | 8.3 | 4.5 $10^{-4}$ | 4.5 $10^{-4}$ - 9 | 0.9983 |

**Table S2: Method validation in Arabidopsis extract.** Validation was conducted according to the protocol described by Hladík et al., 2023. Method precision (expressed as % BIAS) and accuracy (expressed as % RSD) were assessed through a spiking experiment. Arabidopsis seedlings (10 mg, homogenized) were extracted in 1 ml Na-phosphate buffer, and the extracts from five samples were pooled. The pooled extract was divided into 200 µl aliquots, with each aliquot spiked with 5 pmol of stable isotope-labelled standards. Unlabelled standards (1 or 10 pmol) were then added to the aliquots. The samples underwent purification using an in-tip µSPE method, and the concentrations of analytes were measured by HPLC-MS/MS with isotope dilution. Additionally, a separate set of plant extracts was processed without unlabelled standards, allowing for the subtraction of endogenous auxin metabolite levels to calculate recovery rates. Each sample was analysed in five replicates.

| **Analyte** | **1 pmol** | | | **10 pmol** | | |
|---|---|---|---|---|---|---|
| | pmol | BIAS (%) | RSD (%) | pmol | BIAS (%) | RSD (%) |
| **PAA** | 1.2 ± 0.42 | **-15** | **37** | 10.1 ± 0.68 | **-1** | **7** |
| **PAA-Asp** | 1.1 ± 0.11 | **-7** | **10** | 10.2 ± 0.30 | **-2** | **3** |
| **PAA-Glu** | 1.0 ± 0.15 | **5** | **16** | 9.6 ± 0.14 | **4** | **1** |
| **PAA-Val** | 0.9 ± 0.03 | **8** | **3** | 10.6 ± 0.34 | **-6** | **3** |
| **PAA-Leu** | 1.0 ± 0.05 | **1** | **5** | 11.3 ± 0.29 | **-13** | **3** |
| **PAA-Phe** | 1.0 ± 0.05 | **3** | **5** | 10.9 ± 0.33 | **-9** | **3** |
| **PAA-Trp** | 0.9 ± 0.03 | **7** | **3** | 11.1 ± 0.47 | **-11** | **4** |
| **PAA-glc*** | 8.5 ± 2.06 | **15** | **24** | 48.3 ± 1.54 | **3** | **5** |

* PAA-glc was spiked with 10 and 50 pmol

**Table S3: Method validation in pea extract.** Validation was conducted according to the protocol described by Hladík et al., 2023. Method precision (expressed as % BIAS) and accuracy (expressed as % RSD) were assessed through a spiking experiment. Pea seedlings (10 mg, homogenized) were extracted in 1 ml Na-phosphate buffer, and the extracts from five samples were pooled. The pooled extract was divided into 200 µl aliquots, with each aliquot spiked with 5 pmol of stable isotope-labelled standards. Unlabelled standards (1 or 10 pmol) were then added to the aliquots. The samples underwent purification using an in-tip µSPE method, and the concentrations of analytes were measured by HPLC-MS/MS with isotope dilution. Additionally, a separate set of plant extracts was processed without unlabelled standards, allowing for the subtraction of endogenous auxin metabolite levels to calculate recovery rates. Each sample was analysed in five replicates.

| **Analyte** | **1 pmol** | | | **10 pmol** | | |
|---|---|---|---|---|---|---|
| | pmol | BIAS (%) | RSD (%) | pmol | BIAS (%) | RSD (%) |
| **PAA** | 1.2 ± 0.17 | **-17** | **14** | 8.3 ± 0.16 | **17** | **2** |
| **PAA-Asp** | 1.0 ± 0.12 | **-2** | **12** | 9.5 ± 0.39 | **5** | **4** |
| **PAA-Glu** | 1.0 ± 0.07 | **2** | **7** | 9.4 ± 0.29 | **6** | **3** |
| **PAA-Val** | 0.7 ± 0.03 | **29** | **4** | 7.5 ± 0.53 | **25** | **7** |
| **PAA-Leu** | 1.0 ± 0.04 | **-5** | **4** | 10.6 ± 0.62 | **-6** | **6** |
| **PAA-Phe** | 1.0 ± 0.07 | **0** | **7** | 10.6 ± 0.49 | **-6** | **5** |
| **PAA-Trp** | 0.9 ± 0.06 | **12** | **6** | 10.3 ± 0.80 | **-3** | **8** |
| **PAA-glc*** | 8.2 ± 0.40 | **18** | **5** | 36.7 ± 2.32 | **27** | **6** |

* PAA-glc was spiked with 10 and 50 pmol

**Table S4: Data collection and refinement statistics.**

| Enzyme | AtGH3.6 | |
|---|---|---|
| PDB ID | **9FXD** | **9FWD** |
| Ligand | Asp + AMP | AMP |
| Space group | $P6_4$ | $P6_4$ |
| Asymmetric unit | 2 monomers | 2 monomers |
| Unit cell (Å) | | |
| a | 197.9 | 197.0 |
| b | 197.9 | 197.0 |
| c | 65.3 | 65.2 |
| $\alpha$ (°) | 90.0 | 90.0 |
| $\beta$ (°) | 90.0 | 90.0 |
| $\gamma$ (°) | 120.0 | 120.0 |
| Diffraction limits by STARANISO (Å) | 2.13/2.13/1.74 | 2.61/ 2.61/1.93 |
| Resolution (Å)[a] | 98.9–1.74 (1.97-1.74) | 85.3–1.93 (1.97-1.74) |
| Observed reflections | 1474201 (59485) | 1224027 (50777) |
| Unique reflections | 96838 (4844) | 58596 (2932) |
| Completeness spherical (%) | 64.2 (10.3) | 53.5 (8.7) |
| Completeness ellipsoidal (%) | 96.0 (70.3) | 95.4 (76.6) |
| $I/\sigma (I)$ | 11.3 (1.9) | 14.6 (2.0) |
| $R_{sym}$ | 0.161 (1.429) | 0.174 (1.727) |
| $R_{meas}$ | 0.166 (1.490) | 0.179 (1.776) |
| $R_{pim}$ | 0.043 (0.421) | 0.039 (0.411) |
| $CC_{1/2}$[b] | 99.9 (70.5) | 99.8 (63.7) |
| Amino acid residues | 1170 | 1170 |
| Water molecules | 676 | 569 |
| $R_{cryst}$ (%) | 0.1976 | 0.2027 |
| $R_{free}$ (%)[c] | 0.2215 | 0.2402 |
| RMSD bond lengths (Å) | 0.009 | 0.008 |
| RMSD bond angles (°) | 1.06 | 1.01 |
| Mean B value (Å$^2$): | | |
| overall | 33.5 | 44.5 |
| protein chains (A/B) | 32.9/34.1 | 43.1/46.4 |
| water molecules | 34.3 | 40.2 |
| AMP (A/B) | 18.2/21.3 | 28.2/32.3 |
| Asp (A/B) | 33.9/35.0 | -/- |
| Ramachandran statistics (%)[d]: | | |
| favored | 98.9 | 98.4 |
| outliers | 0.0 | 0.0 |
| Molprobity Clashscore[d] | 1.99 | 1.45 |
| Molprobity Overall score[d] | 1.01 | 0.88 |

[a] Numbers in parentheses represent values in the highest resolution shell.

[b] $CC_{1/2}$ stands for a percentage of correlation between intensities from a random half-dataset.

[c] The 5% test set.

[d] Generated with MolProbity (Chen et al 2010).

**Table S5: Ligand Lead Finder rank (LF), docking score and ΔG (Gibbs free energy) score for selected amino acid substrates and products.** Docking was performed in FLARE (https://www.cresset-group.com).

| | AtGH 3.5 | | AtGH 3.6 | |
|---|---|---|---|---|
| **Ligand** | LF Rank score | LF ΔG kcal·mol$^{-1}$ | LF Rank score | LF ΔG kcal·mol$^{-1}$ |
| Asp | −5.315 | −7.560 | −6.615 | −7.101 |
| Glu | −5.164 | −7.285 | −6.669 | −7.611 |
| IAA-Asp | −8.795 | −7.361 | −9.786 | −8.563 |
| IAA-Glu | −7.841 | −6.992 | −9.510 | −7.842 |
| PAA-Asp | −7.107 | −6.800 | −8.560 | −7.530 |
| PAA-Glu | −7.235 | −6.392 | −8.662 | − 7.521 |

### Supplemental experimental procedures: Synthesis of PAA conjugates (Methods S1)

#### Chemicals and general methods

Chemicals and solvents were purchased from common commercial suppliers. All reactions were performed in oven-dried glassware. Conversion of starting materials was monitored by thin layer chromatography (TLC) on aluminium plates coated with silica gel 60 F254 (Merck, USA) and the reaction components were visualised by UV light (254 and 365 nm) and staining solutions (ninhydrin or potassium permanganate). Reaction mixtures were purified by crystallization or column chromatography on silica gel (40-63 µm Davisil LC60A, Grace Davison, UK). $^{1}H$ (500 MHz) and $^{13}C$ (125 MHz) NMR spectra were recorded in deuterated solvents at room temperature on a Jeol ECA-500 spectrometer equipped with a 5 mm Royal probe and compared with reported data. The LC–MS analyses were performed on an ACQUITY UPLC® H-Class system combined with UPLC® PDA detector and a single-quadrupole mass spectrometer QDa™ (Waters, UK) as described previously (Bieleszová et al., 2019).

#### Synthesis of PAA-NHS and [$^{13}C_6$]PAA-NHS

Phenylacetic acid (1950 mg, 14.3 mmol) was dissolved in dioxane/ethyl acetate (15/7; 22 mL), cooled down to 0°C and *N*-hydroxysuccinimide (1730 mg, 15 mmol) and *N,N´*-dicyclohexylcarbodiimide (3090 mg, 15 mmol) were added sequentially. Reaction mixture was brought up to room temperature and stirred for 1 h. Upon completion, reaction mixture was cooled down to 0°C, fileted through a pad of Celite and the filter cake was washed with dioxane/ethyl acetate (3/1; 2 × 15 mL). The filtrate was evaporated to dryness and the residue was purified by crystallization from 2-propanol (30 mL) to give pure PAA-NHS (3010 mg; 90%), the spectral data of which was in good agreement with the published data (Zasedateleva et al., 2020). [$^{13}C_6$]PAA-NHS was prepared analogously using [$^{13}C_6$]-phenylacetic acid as a starting material.

#### Synthesis of PAA-Asp and [$^{13}C_6$]PAA-Asp

PAA-NHS (1000 mg, 4.25mmol) was reacted with L-aspartic acid sodium salt (769 mg, 4.96 mmol) in dioxane/water (1/1; 80 mL) at room temperature for 3 h. Upon completion, reaction mixture was cooled down to 0°C, acidified with 1M HCl to pH = 2.5-3 and extracted with ethyl acetate (3 × 40 mL). Combined organic extracts were washed with brine (40 mL), dried over sodium sulphate and evaporated to dryness. The residue was purified by column chromatography to give pure PAA-Asp (570 mg; 53%), the spectral data of which was in good agreement with the published data (Thangavelu et al., 2017). [$^{13}C_6$]PAA-Asp was prepared analogously using [$^{13}C_6$]PAA-NHS as a starting material.

#### Synthesis of PAA-Glu and [$^{13}C_6$]PAA-Glu

PAA-NHS (500 mg, 2.14 mmol) was reacted with L-glutamic acid sodium salt (419 mg, 2.48 mmol) in dioxane/water (1/1; 40 mL) at room temperature for 3 h. Upon completion, reaction mixture was cooled down to 0°C, acidified with 1M HCl to pH = 2.5-3 and extracted with ethyl acetate (3 × 20 mL). Combined organic extracts were washed with brine (20 mL), dried over sodium sulphate and evaporated to dryness. The residue was purified by column chromatography to give pure PAA-Glu (195 mg; 34%), the spectral data of which was in good agreement with the published data (Thangavelu et al., 2017). [$^{13}C_6$]PAA-Glu was prepared analogously using [$^{13}C_6$]PAA-NHS as a starting material.

#### Synthesis of PAA-Val

PAA-NHS (1500 mg, 6.4 mmol) was reacted with L-valine sodium salt (1043 mg, 7.5 mmol) in dioxane/water (1/1; 100 mL) at room temperature for 3 h. Upon completion, reaction mixture was cooled down to 0°C, acidified with 1M HCl to pH = 2.5-3 and extracted with ethyl acetate (3 × 50 mL). Combined organic extracts were washed with brine (50 mL), dried over sodium sulphate and evaporated to dryness. The residue was purified by column chromatography to give pure PAA-Val (1400 mg; 92%), the spectral data of which was in good agreement with the published data (Schwieter and Johnston, 2016).

**Synthesis of PAA-Phe**

PAA-NHS (1500 mg, 6.4 mmol) was reacted with L-phenylalanine sodium salt (1404 mg, 7.5 mmol) in dioxane/water (1/1; 100 mL) at room temperature for 3 h. Upon completion, reaction mixture was cooled down to 0°C, acidified with 1M HCl to pH = 2.5-3 and extracted with ethyl acetate (3 × 50 mL). Combined organic extracts were washed with brine (50 mL), dried over sodium sulphate and evaporated to dryness. The residue was purified by column chromatography to give pure PAA-Phe (1311 mg; 72%), the spectral data of which was in good agreement with the published data (Schwieter and Johnston, 2016).

**Synthesis of PAA-Leu**

PAA-NHS (1500 mg, 6.4 mmol) was reacted with L-leucine sodium salt (1149 mg, 7.5 mmol) in dioxane/water (1/1; 100 mL) at room temperature for 3 h. Upon completion, reaction mixture was cooled down to 0°C, acidified with 1M HCl to pH = 2.5-3 and extracted with ethyl acetate (3 × 50 mL). Combined organic extracts were washed with brine (50 mL), dried over sodium sulphate and evaporated to dryness. The residue was purified by column chromatography to give pure PAA-Leu (1395 mg; 87%), the spectral data of which was in good agreement with the published data (Gabor and Janssen, 2004).

**Synthesis of PAA-Trp**

PAA-NHS (1500 mg, 6.4 mmol) was reacted with L-tryptophan sodium salt 1697 mg, 7.5 mmol) in dioxane/water (1/1; 100 mL) at room temperature for 3 h. Upon completion, reaction mixture was cooled down to 0°C, acidified with 1M HCl to pH = 2.5-3 and extracted with ethyl acetate (3 × 50 mL). Combined organic extracts were washed with brine (50 mL), dried over sodium sulphate and evaporated to dryness. The residue was purified by column chromatography to give pure PAA-Trp (1120 mg; 54%), the spectral data of which was in good agreement with the published data (Matsui et al., 2021).

**Synthesis of PAA-glc**

Glucose (199 mg, 1.1 mmol) was brought up into anhydrous dioxane (37 mL) and sonicated under argon for 15 min. Subsequently, phenylacetic acid (50 mg, 0.37 mmol) and triphenylphosphine (193 mg, 0.74 mmol) were added to the reaction mixture, followed by a dropwise addition of diisopropyl azodicarboxylate (144 µL, 0.74 mmol). Resulting mixture was stirred vigorously at room temperature for 30 min, quenched with methanol, and evaporated to dryness. The residue was purified by column chromatography to give pure PAA-glc (52 mg; 47%), the spectral data of which was in good agreement with the published data (Iddon et al., 2011).
